# Phytoplankton consortia as a blueprint for mutually beneficial eukaryote-bacteria ecosystems: Biocoenosis of *Botryococcus* consortia

**DOI:** 10.1101/476887

**Authors:** Olga Blifernez-Klassen, Viktor Klassen, Daniel Wibberg, Enis Cebeci, Christian Henke, Christian Rückert, Swapnil Chaudhari, Oliver Rupp, Jochen Blom, Anika Winkler, Arwa Al-Dilaimi, Alexander Goesmann, Alexander Sczyrba, Jörn Kalinowski, Andrea Bräutigam, Olaf Kruse

## Abstract

Bacteria occupy all major ecosystems and maintain an intensive relationship to the eukaryotes, developing together into complex biomes (i.e., phycosphere and rhizosphere). Interactions between eukaryotes and bacteria range from cooperative to competitive, with the associated microorganisms affecting their host’s development, growth, health and disease. Since the advent of non-culture dependent analytical techniques such as metagenome sequencing, consortia have been described but owing to the complex interactions rarely functionally dissected. Multifaceted analysis of the microbial consortium of the ancient phytoplankton *Botryococcus* as an attractive model food web revealed that its all abundant bacterial members belong to a distinct niche of biotin auxotrophs, essentially depending on the microalga. In addition, hydrocarbonoclastic bacteria without vitamin auxotrophies, which adversely affect the algal cell morphology, appear evidently decimated. Synthetic rearrangement of a minimal community consisting of alga, mutualistic and parasitic bacteria underpins the model of a eukaryote that domesticates its own mutualistic bacterial “zoo” to manipulate and control its surrounding biosphere. This model of domestication of mutualistic bacteria for the defense against destruents by a eukaryotic host could represent ecologically relevant interactions that cross species boundaries. Metabolic and system reconstruction disentangles the relationships and provide a blueprint for the construction of mutually beneficial synthetic ecosystems.

## Background

Bacteria are omnipresent and affect likely every eukaryotic system through their presence (Grosberg and Strathmann, 2007). Microalgae are the dominant primary producers and the base of the food web within aquatic ecosystems and sustain in the natural environment in close relationship with multiple associated microorganisms (Ramanan et al., 2016; Seymour et al., 2017). Thus, the interactions between phytoplankton and bacteria represent a fundamental ecological relationship within aquatic environments (Seymour et al., 2017), by controlling nutrient cycling and biomass production at the food web base. The nature of the exchange of micro- and macronutrients, diverse metabolites including complex products such as polysaccharides, and infochemicals defines the relationship of the symbiotic partners (Cole, 1982; Amin et al., 2015; Zengler and Zaramela, 2018), which span mutualism, commensalism and parasitism (Ramanan et al., 2016). Within a parasitic association, the bacteria are known to adversely affect the algae (Wang et al., 2010), while in commensalism the partners can survive independently, with the commensal feeding on algae-provided substrates, but without harming the host (Ramanan et al., 2016). Mutualistic interactions include *i.a.* the obligate relationships between vitamin-synthesizing bacteria and vitamin-auxotrophic phytoplankton species (Croft et al., 2005). Many microalgae cannot synthesize several growth-essential vitamins (like B_12_ and B_1_) (Croft et al., 2005; Croft et al., 2006; Tang et al., 2010) and obtain them through a symbiotic relationship with bacteria in exchange for organic carbon (Grant et al., 2014). Recent observations suggest that not only certain microalgae require vitamins, but many bacteria also have to compete with other organisms for the B-vitamins (Sañudo-Wilhelmy et al., 2014; Gómez-Consarnau et al., 2018).

However, in microbial environments, where the distinction between host and symbionts is less clear, the identification of the associated partners and the nature of their relationship is challenging. Looking for a suitable model food web system to elucidate the complexity of eukaryote-prokaryote interactions, we focused on the microalga-bacteria consortia with an organic carbon-rich phycosphere. The planktonic chlorophyta *Botryococcus braunii* excretes chemically inert long-chain hydrocarbons and a variety of exo-polysaccharides (Banerjee et al., 2002) which allows this single celled alga to form large agglomerates (Grosberg and Strathmann, 2007). Both organic carbon-based products are exposed to the surrounding ecosystem and enable the alga to sustain in the natural environment (phycosphere) in close relationship with multiple associated microorganisms (Ramanan et al., 2016; Seymour et al., 2017). *Botryococcus* exudes up to 46% of their photosynthetically fixed carbon into the phycosphere (Blifernez-Klassen et al., 2018), similarly to processes, however with lesser extent, observed for seedlings as well as adult plants within the rhizosphere in the form of poorly characterized rhizodeposits (Nguyen, 2003; Bisseling et al., 2009; Marschner, 2011). *B. braunii* consortia, naturally accompanied with a variety of microorganisms (Chirac et al., 1985), may represent an example for an association already existing during the early stages of life development on earth, since it is generally considered that a progenitor of *Botryococcus* impacted the development of today’s oil reserves and coal shale deposits (McKirdy et al., 1986). Thus, the microbiome associated with *Botryococcus braunii* presents a unique opportunity to dissect the roles of the eukaryote and prokaryotes within the consortium and test if and how the eukaryote manipulates its biome.

The interactions between *B. braunii* and the associated bacteria are not fully understood, but are known to influence the microalgal growth performance and product formation capacity (Chirac et al., 1985; Banerjee et al., 2002; Tanabe et al., 2015; Fuentes et al., 2016). To understand the complex nature of the unique habitat of a microalgal phycosphere and to unravel the food web between alga and bacteria, we investigated the consortia accompanying the hydrocarbon/carbohydrate-producing green microalga *Botryococcus braunii*.

## Results and discussion

### Metagenomic survey of the *Botryococcus braunii* consortia

To determine the inhabitants of the *B. braunii* ecosystem, we profiled four different *Botryococcus* communities (races A and B) using high-throughput 16S rDNA gene amplicon and read-based metagenome sequencing approaches (Figures S1-3; Figure 1a). Assembly and functional annotation of the metagenomes together with single strain isolation and sequencing reconstructed the capabilities of the most abundant consortial microorganisms and were used to design and test synthetic alga-bacteria ecosystems. The analysis of the 16S rDNA amplicons and a read-based metagenome approach showed that each of the *B. braunii* communities represents a comparable conserved consortium of a single algal and various bacterial species as well as traces of Archaea and viruses (Figure S2). 16S rDNA amplicon sequencing identified 33 bacterial taxa (Figure S3). The relative abundance of plastid 16S rDNA sequences accounted up to 49 and 59% of all detected amplicon reads in race A and B samples, respectively. Metagenomic assembly resulted in ten high quality draft genomes with >80% assembled. Three additional genomes were closed by single strain sequencing (Figure 1; Tables S1 and S2; Supplementary Discussion, Chapter 1). The bacterial community varied quantitatively, but not qualitatively between growth phases and strains (Figure S3). *B. braunii* consortia are inhabited by the same bacterial phyla (*Proteobactreria, Bacteroidetes, Acidobacteria* and *Actinobacteria*), albeit in different relative abundances, compared to those phyla which occupy the plant rhizosphere (Xu et al., 2018) and the mammals gut (Ley et al., 2008).

**Figure 1:**
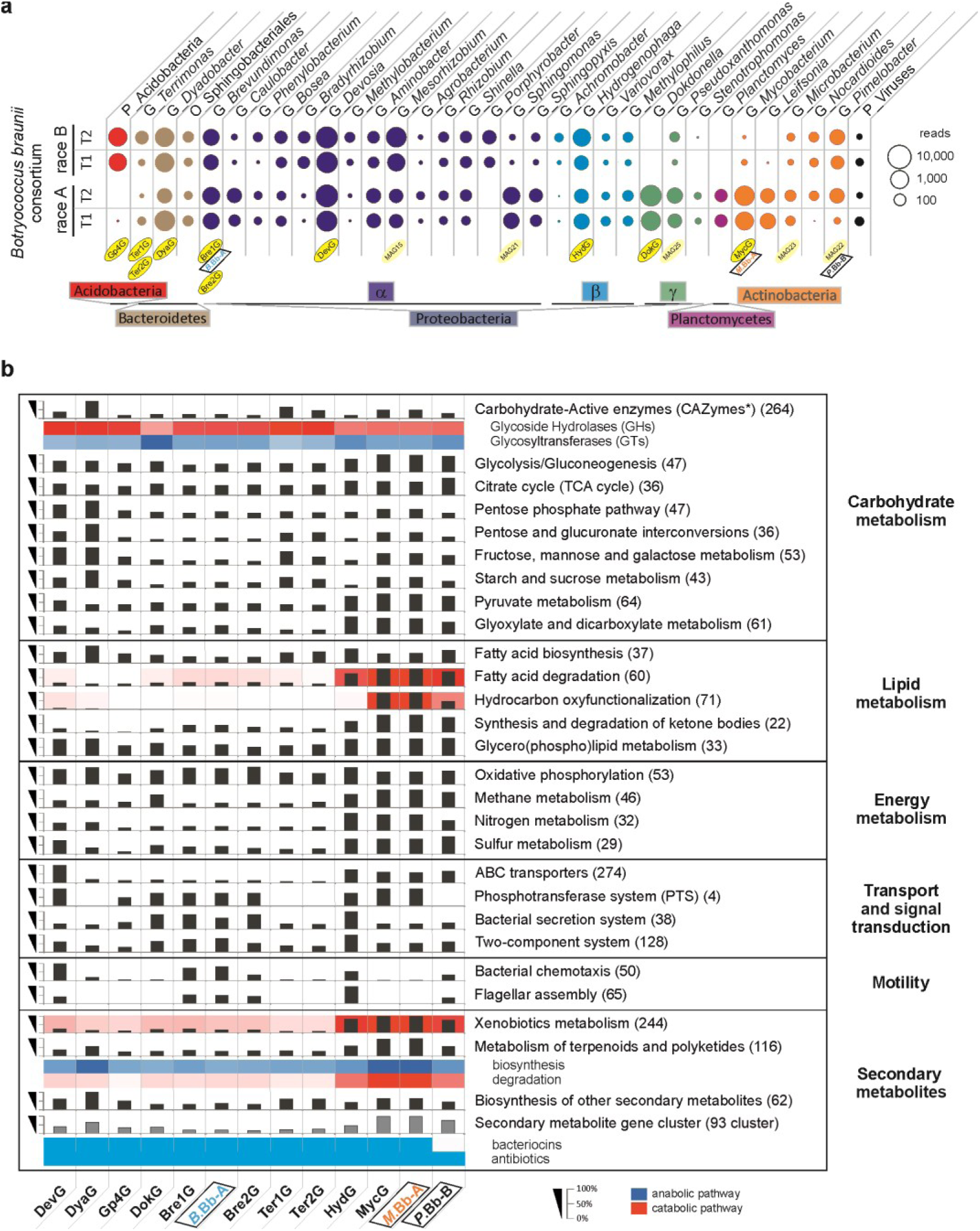
Taxonomy of the *B. braunii* bacterial community and selected functional categories encoded in reconstructed MAGs and complete genomes. (**a**) Normalized read-based comparison and taxonomic assignment of different *B. braunii* metagenome datasets. The circle size represents the amount of classified reads for the respective taxonomic group (minimum 50 reads per taxonomic group). (**b**) Abundance of selected functional categories encoded in the high-quality draft and complete bacterial genomes. Numbers in parentheses represent the maximal total number of genes within each pathway. Percentage: per cent of total number of genes per pathway identified within each genome. All annotated genes within the respective pathways are presented on the EMGB platform (https://emgb.cebitec.uni-bielefeld.de/Bbraunii-bacterial-consortium/). The pathways statistics are summarized in the Table S5.

During the growth phase, the *Botryococcus* phycosphere is dominated by the genera of *Devosia, Dyadobacter, Dokdonella, Acidobacteria, Brevundimonas, Mesorhizobium* and *Hydrogenophaga* which account for over 80% of the bacteria (Table S1). The re-processing of the *B. braunii* Guadeloupe strain metagenome data (Sambles et al., 2017) and direct comparison of the different datasets revealed the coinciding presence of the individual bacterial genera, although massively varying in the abundance pattern in dependence of cultivation conditions (Figure S5).The same taxa are also present *i.a.* in the active microbiome associated roots and rhizosphere of oilseed rape (Gkarmiri et al., 2017), as well as in the lettuce (Schreiter et al., 2014) and global citrus rhizosphere (Xu et al., 2018), strongly suggesting an overall mutual dependence or preferred coexistence.

To identify genes that enable the bacterial community members to contribute to and to benefit from the food web of the consortium, we performed a KEGG-based quantitative functional assignment of the annotated high-quality MAGs (metagenome assembled genomes). Based on these data sets, we sequenced genomes of isolated associates and imported them to the EMGB platform (elastic metagenome browser, https://emgb.cebitec.uni-bielefeld.de/Bbraunii-bacterial-consortium/; for details see Methods; Supplementary Discussion, Chapter 2). We originally expected a uniform distribution of abilities and a high degree of interconnectedness between all members of the consortium. Surprisingly, the consortium divides into two functionally distinct groups of bacteria: The dominant consortia members such as *Devosia* (DevG), *Dyadobacter* (DyaG) and *Brevundimonas* (Bre1G, *B.*Bb-A) do not oxyfunctionalize hydrocarbons (Rojo, 2010), but their genomes are rich in genes involved in the degradation/assimilation of carbohydrates (Figure 1b). On the contrary, the bacterial strains *Mycobacterium* and *Pimelobacter* (*M*.Bb-A and *P*.Bb-B), which are of low abundance within the consortia (Figure S3), contain a large portfolio of genes encoding monooxygenases/hydroxylases for hydrocarbon degradation (Rojo, 2010) and hold the potential for a pronounced catabolic capacity towards lipids, fatty acids and xenobiotics (Figure 1b). Besides, *Mycobacterium* and *Pimelobacter* genomes were the only genomes among the abundant community members to carry non-heme membrane-associated monooxygenase *alkB* genes (Table S6). To test these observed genetic predisposition of the consortia, we assessed algal capacities for growth and hydrocarbon production in co-cultivation of the axenic *B. braunii* with one representative of each group.

### Effect of the bacterial community on the *B. braunii* host

One representative of the core as well as less abundant community members were isolated in axenic culture (*Brevundimonas* sp. Bb-A and *Mycobacterium* sp. Bb-A, respectively). Synthetic algae-bacteria consortia were formed with an axenic hydrocarbon- and carbohydrate producing *B. braunii* race A strain under strict phototrophic conditions (*e.g.* without the supplementation of vitamins or organic carbon). Axenic *Botryococcus braunii* showed a continuous increase in cell biomass and hydrocarbon content, yielding 2.1±0.2 gL^-1^ and 0.7±0.1 gL^-1^, respectively (Figure 2a, b; Figure S6). Co-cultivation of the *Brevundimonas* sp. Bb-A promotes growth of the alga and increases biomass by 60.5±16.9% compared to axenic controls (Figure 2a) while co-cultivation with the *Mycobacterium* sp. Bb-A reduces growth by 30.6±4.8% (Figure 2b).

**Figure 2:**
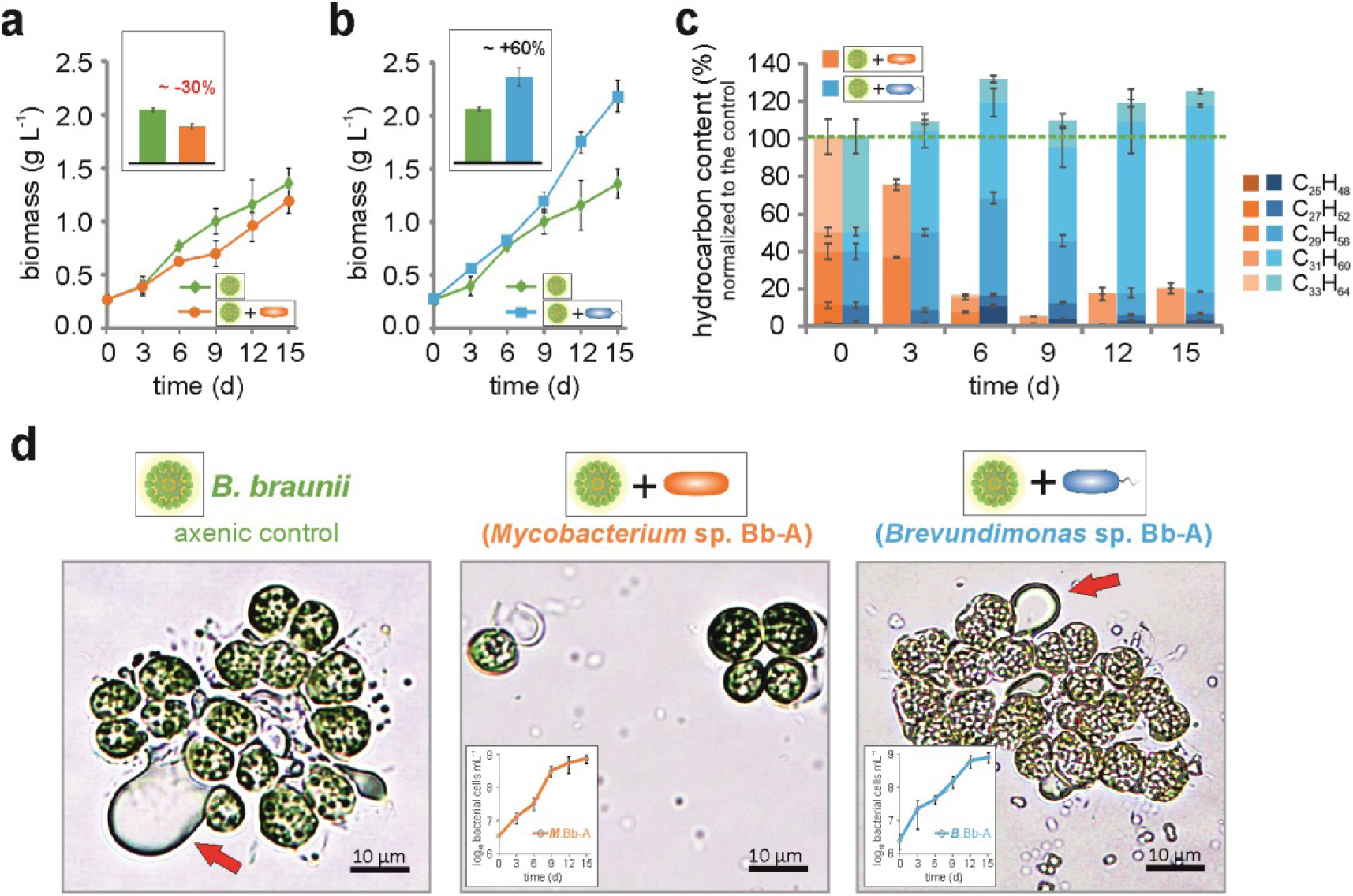
Physiological effect of the individual bacterial isolates on the microalgae during co-cultivation. (**a**-**d**) Determination of algal growth and product formation performance of axenic and xenic *B. braunii* cultures, supplemented with bacterial isolates. Shown are growth analysis of *B. braunii* in the presence of (**a**) *Mycobacterium* sp. Bb-A and (**b**) *Brevundimonas* sp. Bb-A; (**c**) comparison of the hydrocarbon levels observed in samples cultivated with bacterial isolates in relation to the axenic *B. braunii* culture in the course of the cultivation; (**d**) morphological characteristic of algal cells during the cultivation (9 days) with the bacterial isolates (red arrows indicate the hydrocarbons produced by the microalga). Graphs imbedded in the pictures show the progress of the bacterial growth (cell number on log_10_ scale).The error bars represent standard error of mean values of three biological and three technical replicates (SE; n=9).

*B. braunii* accumulates hydrocarbons up to 38% of its dry weight (DW) in a bacteria-free environment. In contrast, co-cultivation with *Mycobacterium* limits hydrocarbon accumulation down to 4.4±0.2% of DW (Figure 2c) and alters colony morphology (Figure 2d). Loss of the hydrocarbon matrix reduces colony size (Figure 2d) and likely increases susceptibility to grazers (Pančić and Kiørboe, 2018). The malfunction of colony formation can thus be attributed to imbalanced and dysbiotic host-microbiome interaction similar to imbalanced interactions previously observed (Gilbert et al., 2016; Durán et al., 2018) and *Mycobacterium* essentially acts as a pathogen. The dramatic decrease in hydrocarbon content and disturbed colony formation cannot be observed in cultures supplemented with *Brevundimonas* or in the xenic *Botryococcus braunii* algae-bacteria communities (Figure 2c, d; Figures S1 and S6), pointing to a balanced equilibrium between the alga and its microbiome in natural environments.

Since *Botryococcus* readily releases huge amounts of organic carbon into the extracellular milieu (Blifernez-Klassen et al., 2018), creating a phycosphere that naturally attracts many microorganisms, including those with potential damaging effects, its evolutionary fitness will depend on efficient management of the “bacterial zoo”. We hypothesized that there is a strong control mechanism that allows the algae host to attract and control symbiotic / beneficial bacteria. Therefore, we examined the genetic portfolio of the bacterial community for essential cofactors and interdependencies.

### The core community of *Botryococcus* phycosphere belongs to a distinct niche of biotin auxotrophs

At first glance, the *Botryococcus* alga does not appear to be obligatory dependent on a bacterial community, since axenic strains can be fully photoautotrophically cultured without vitamin supplements (Figure 2 a, b) and contains the vitamin B_12_-independent form of methionine synthase (*metE*) (Molnár et al., 2012). (Meta)Genome reconstruction revealed that the *B. braunii* bacterial core community is auxotrophic for various B-vitamins (Figure 3a), making them dependent on an exogenous supply of vitamins for survival. All abundant genera within the *B. braunii* consortia (including the growth-promoting *Brevundimonas*) lack the complete gene portfolio for biotin (vitamin B_7_) synthesis (Figure 3a), but contained the complete biosynthesis pathway of fatty acids for which biotin is essential (Sañudo-Wilhelmy et al., 2014). The complete thiamin synthesis pathway (vitamin B_1,_ essential for citrate cycle and pentose phosphate pathway (Sañudo-Wilhelmy et al., 2014)) could not be reconstructed for four assembled genomes. Five genera are auxotrophic for cobalamin (vitamin B_12_). Although, auxotrophic for various B-vitamins including biotin, the consortia’s most abundant species *Devosia* (DevG) contains the complete pathway for vitamin B_12_ synthesis. In view of the fact that the bacterial abundance determines their functional relevance for the community (Reed and Martiny, 2007; Rivett and Bell, 2018), this species is also likely to provide both the bacterial community and the microalga with this vitamin (Croft et al., 2005), thus confirming the recent proposition that the B_12_ supply is realized only by a few certain bacteria within the alga-bacteria consortia (Krohn-Molt et al., 2017). In this context, since the B_7_-auxotrophs are prevalent within the detected consortia, it is doubtful that the large required supply of biotin is of bacterial origin. Because of its predominance within the community (Figure S2), the microalga is more likely to secrete and provide this vitamin to the bacteria.

**Figure 3:**
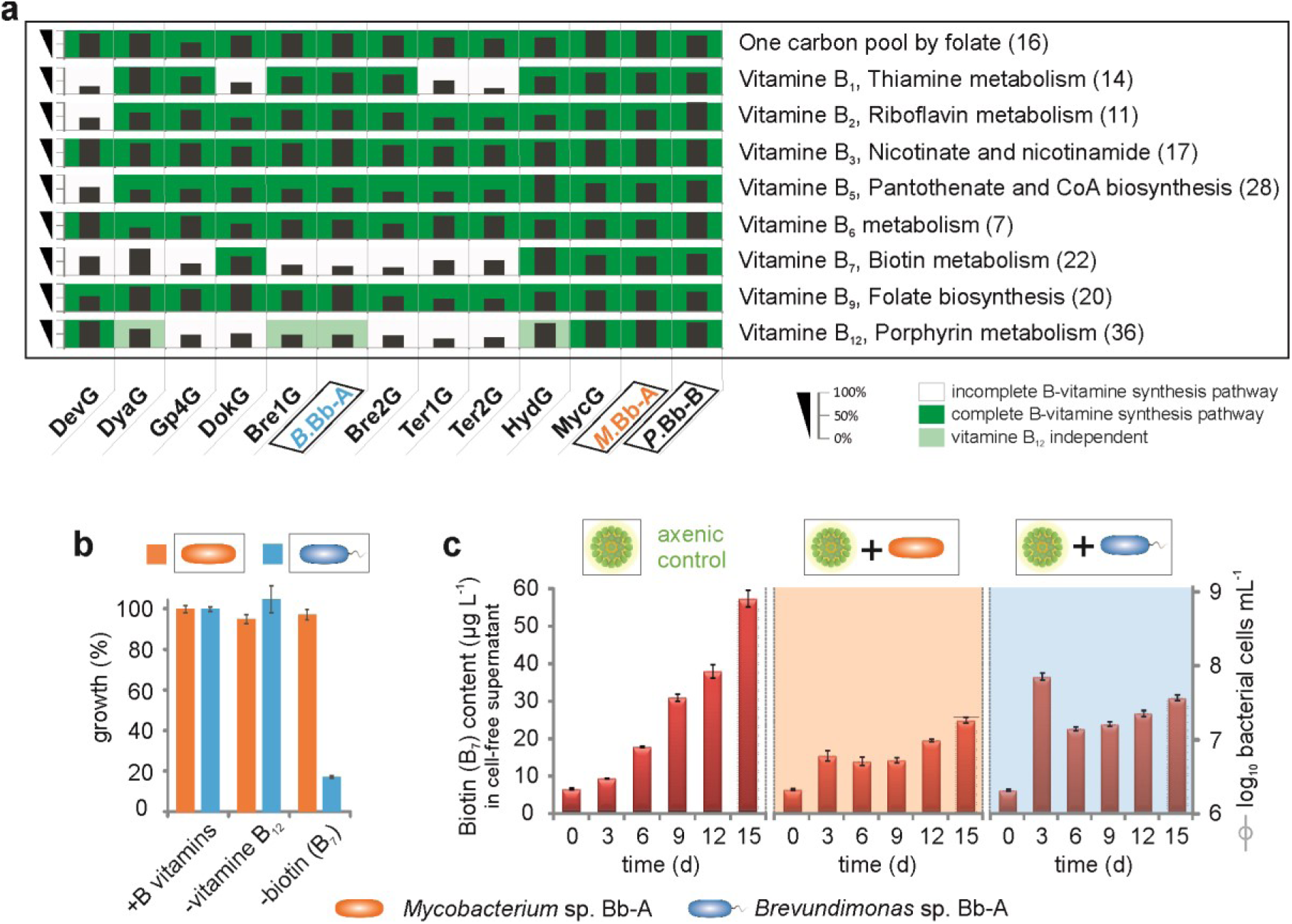
Essential cofactors (inter-)dependencies of the bacterial core community and isolates of *B. braunii*. (**a**) Heatmap of the reconstructed *de novo* biosynthesis of B-vitamins encoded in the high-quality draft and complete bacterial genomes. Numbers in parentheses represent the maximal total number of genes within each pathway. Percentage: per cent of total number of genes per pathway identified within each genome. All annotated genes within the respective pathways are presented on the EMGB platform (https://emgb.cebitec.uni-bielefeld.de/Bbraunii-bacterial-consortium/). The pathways statistics are summarized in the Table S5 and S8. (**b**) Growth assay to confirm B-vitamin auxotrophy and prototrophy in axenic *Brevundimonas* sp. Bb-A (blue) and *Mycobacterium* sp. Bb-A (orange) strains, respectively. (**c**) Biotin concentration levels detected in cell-free supernatants from the cultures of axenic *B. braunii* as well as in the presence of *Mycobacterium* sp. Bb-A (orange) and *Brevundimonas* sp. Bb-A (blue). The error bars represent standard error of mean values of three biological and three technical replicates (SE; n=9).

It is well known, that many microalgae rely on the external supply of growth-essential vitamins like cobalamin and thiamin (Croft et al., 2005; Croft et al., 2006; Tang et al., 2010), however, biotin auxotrophy is rare among phytoplankton species (Croft et al., 2006). Here, our genome reconstruction results strongly suggests that also bacteria have to compete with other organisms for the B-vitamins and live in a close mutualistic relationship with the microalga strongly depending on the biotin supply. The growth promoting effect of *Brevundimonas* (Figure 2b), which depends on an exogenous supply of biotin for survival, is thus based on the close interaction with the microalga. In contrast, low abundant consortial members such as *Mycobacterium* are B-vitamin prototrophs (Figure 3a).

Extended experimental analysis confirmed the biotin (vitamin B_7_) auxotrophy in *Brevundimonas* sp. Bb-A and vitamin B prototrophy in *Mycobacterium* sp. Bb-A as deduced from the assembled genomes (Figure 3b). *Botryococcus braunii* secrets biotin into the surrounding environment (Figure 3c) and secretion is increasingly induced by the presence of bacteria, particularly in the cultures with the B_7_-auxotrophs (compare to Figure 3c at timepoint 3 days). The secretion of some B-vitamins has already been observed in microalgae (Aaronson et al., 1977), and also plants support their rhizosphere by active riboflavin (vitamin B_2_) excretion from their roots (Higa et al., 2008). However, the eukaryotic B-vitamin secretion (especially of biotin), which was increasingly induced by the presence of bacteria, as well as their simultaneous active uptake, remained so far unrecognized.

Collectively, in a healthy (natural) community, non-hydrocarbon degrading bacterial genera which are auxotrophic for one or more vitamins dominate the phycosphere, while fully prototrophic hydrocarbonoclastic bacteria are present but not abundant. We hypothesized that *B. braunii* cultivates its own auxotrophic “bacterial zoo” to combat hydrocarbonoclastic bacteria. To test the hypothesis, synthetic consortia of alga–”good bacteria”–”bad bacteria” were formed and analyzed.

### Bacterial biotin-auxotrophic symbionts fend their eukaryotic host

We hypothesized that the alga maintains its balanced, ‘healthy’ microbiome by recruiting beneficial, vitamin B-auxotrophic microorganisms with the provision of photosynthetically fixed carbon and biotin and thus counteracts pathogen assault. *Brevundimonas* and *Mycobacterium* were co-cultivated with the axenic *Botryococcus* alga under photoautrophic conditions with three different inoculum ratios (Figure 4a).

**Figure 4:**
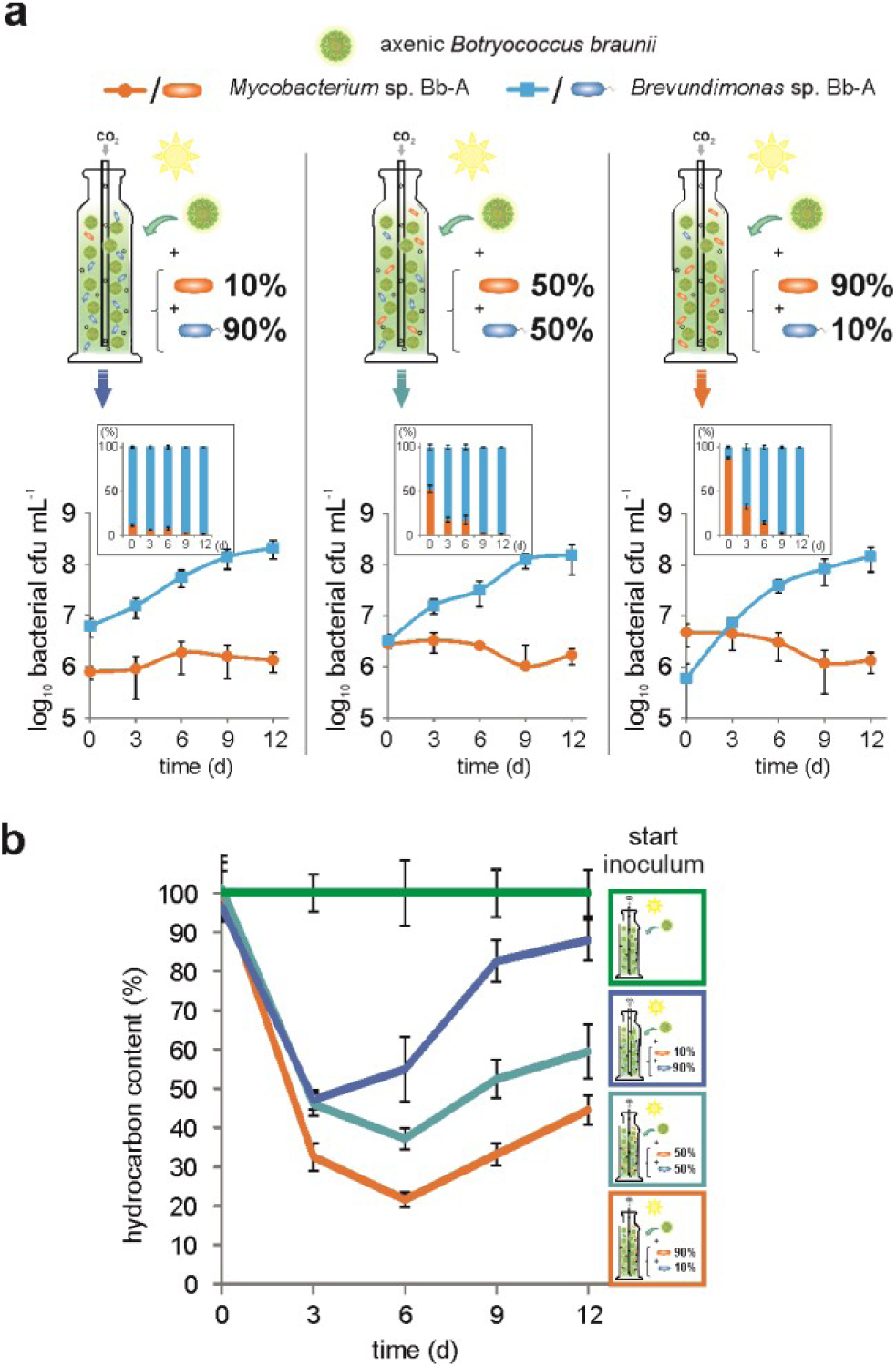
Bacterial symbionts fend their eukaryotic host from hydrocarbonoclastic invaders. Physiology of a synthetic alga-bacteria-bacteria community, consisting of *Brevundimonas* sp. Bb-A (blue) and *Mycobacterium* sp. Bb-A (orange) in co-culture with *B. braunii* (green) under photoautrophic conditions with three different inoculum ratios. Shown are (**a**) the progression of bacterial proliferation (cfu, colony-forming units) and (**b**) detected relative hydrocarbon levels (normalized to the axenic control) during the co-cultivation. The error bars represent standard error of mean values of three biological and three technical replicates (SE; n=9).

Independently from the initial amount of inoculum, biotin-auxotrophic *Brevundimonas* proliferated strongly within this synthetic community. *Mycobacterium* did not proliferate in the presence of *Brevundimonas* (Figure 4a), but was clearly capable of growth in the presence of the alga alone (Figure 2b). *Brevundimonas* apparently produces bioactive compounds with bacteriostatic properties, which likely decimate the proliferation of *Mycobacterium*. Indeed, all reconstructed genomes encode metabolic gene cluster for antibiotic compounds (especially against Gram-positive bacteria) and for bacteriocins (Figure 1b, Table S9) with significant antimicrobial potency (Cotter et al., 2013), the potential of which should be subject of further research. The protective effect of *Brevundimonas* is evident from the hydrocarbon content of cultures which increases with decreasing proportions of *Mycobacterium* (Figure 4b). Experimental analyses (Figures 2-4) thus confirmed key conjectures derived from metagenomic assemblies (Figure 1).

In conclusion, the interactions between the microalga and bacteria within the *Botryococcus braunii* consortia depends on a complex interplay between all members, with the microalga as the “conductor” in this play (Figure 5). Our observations indicate that the microalga is associated with a “zoo” of largely supportive bacteria and suggest a framework in which the microalga specifically attracts vitamin-auxotrophic bacterial species and domesticates them by controlling vitamin supply. Since autark bacteria without vitamin auxotrophies are present in small proportions only (Figure S3; Figure 3a), they are likely decimated by antibacterial agents from the supportive community.

**Figure 5:**
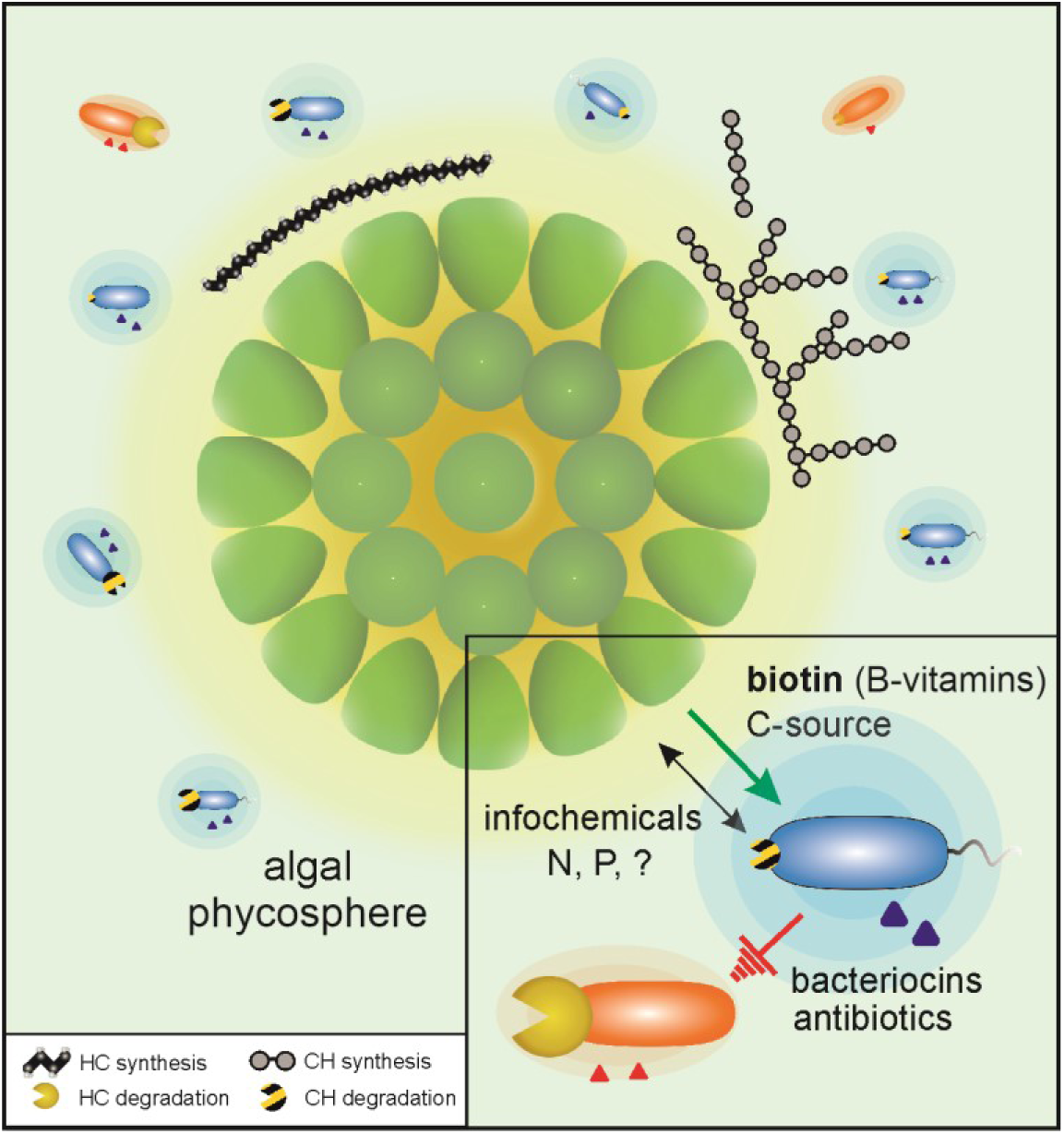
Alga-bacteria interactions within the *Botryococcus braunii* phycosphere. The depiction summarizes the observed generic potential encoded in the (meta)genomes of bacterial associates (Figures 1 and 3), along with experimental validation within the minimal synthesis community (Figures 2 and 4). The color code indicates the functional grouping of the community members: supportive B-vitamin-auxotrophic (blue) and hydrocarbon-clastic prototrophic (red). The potential for degradation of hydrocarbons and carbohydrates of the bacterial associates is illustrated via yellow and striped pacmans, respectively. The genetic predisposition to the synthesis of bacteriocins and antibiotics encoded by the secondary metabolite biosynthesis gene clusters, is indicated by the halos and triangles, respectively. Abbreviation: HC, Hydrocarbons; CH, carbohydrates; B_7_, Biotin; N, nitrogen; P, phosphorus.

The analyses mechanistically resolved biotin as a critical metabolite and indicated the presence of at least two additional communication signals: a bacterial signal increasing biotin production (Figure 3c) and a bacteriostatic compound directed from *Brevundimonas* at *Mycobacterium* (Figure 4a). The metagenomes hint at additional beneficial interactions between the alga and its cultivated “zoo”: vitamin B_12_, and assimilated N and P from the bacteria in exchange for biotin (and other vitamins) as well as carbon from the alga (Figure 5). The successful design of minimal synthetic alga-bacteria cultures consisting only of two bacteria species and the alga precisely demonstrates a functional triangle relationship for preventing uncontrolled overgrowing of hydrocarbonoclastic destruents. However, the elucidation of the communication signals exchanged between the participating parties will be the subject of further research.

The metagenomic analysis of the *Botryococcus* consortium presents a path towards mechanistic resolution of ecological interactions at the microscopic scale. Metabolic and system reconstruction unravels the relations in the eukaryote-bacteria consortia and provides a blueprint for the construction of mutually beneficial synthetic ecosystems.

## Methods

### Strains, cultivation conditions and growth determination parameters

Liquid, xenic cultures of *Botryococcus braunii* race A (CCALA778, isolated in Serra da Estrela, Baragem da Erva da Fome, Portugal) and race B (AC761, isolated from Paquemar, Martinique, France) as well as the axenic *B. braunii* race A strain (SAG30.81, Peru, Dpto.Cuzco, Laguna Huaypo) were used for co-cultivation studies. The cultures were phototrophically cultivated under 16:8 light-dark illumination of 350±400 μmol photons m^-2^ s^-1^ white light in modified Chu-13 media (without the addition of vitamin and organic carbon source) as described before (Blifernez-Klassen et al., 2018). Carbon supply and agitation were achieved by bubbling the cultures with moisture pre-saturated carbon dioxide-enriched air (5%, v/v) with a gas flow rate of 0.05 vvm. Cultures growth was monitored by measurements of biomass organic dry weight (DW) by drying 10 mL culture per sample (in three biological and technical replicates) following the protocol described before (Astals et al., 2012). Additionally, microalgae and bacteria cells were regularly checked by optical microscopy.

The bacterial isolates were obtained by plating highly diluted xenic *B. braunii* cultures (CCALA778 and AC761) on LB agar plates supplied with vitamins (0.1 nM cobalamin, 3.26 nM thiamine and 0.1 nM biotin). After 1-2 weeks single cell colonies were picked, separately plated and analyzed *via* 16S rDNA Sanger-sequencing by using the universal primer pair 27F (5’-AGAGTTTGATCCTGGCTCAG-3’) and 1385R (5’-CGGTGTGTRCAAGGCCC-3’). Two isolates, deriving from algal race A and one from race B, were classified based on full length 16S rDNA similarity as *Mycobacterium* sp., *Brevundimonas* sp. and *Pimelobacter* sp., respectively. *Mycobacterium* sp. and *Brevundimonas* sp. were used for further analysis. The purity of the axenic alga and bacterial strains was regularly verified by polymerase chain reaction (PCR) amplification and Sanger-sequencing of the 16S rRNA gene using a universal primer pair 27F and 1385R.

For experiments testing the requirement of B-vitamins, mono-cultures of *Mycobacterium* sp. Bb-A and *Brevundimonas* sp. Bb-A were pre-washed three times in minimal medium (Schlegel et al., 1961) and then starved of vitamins for two days prior to the start of the experiment. Then, bacterial cells were diluted to 0.2 (oD_600_) with minimal medium supplemented with 1gL^-1^ glucose and B-vitamins (at the following concentrations: 0.1 nM (each) of cobalamin, riboflavin, biotin and pantothenate as well as 3.26 nM thiamine) and grown for seven days.

For the co-cultivation experiments, 500 mL axenic algal cultures (∼0.25g L^-1^) were inoculated with the isolates (*Mycobacterium* sp. Bb-A or *Brevundimonas* sp. Bb-A, with ∼2×10^6^ cell mL^-1^ for single isolate cultivation and with ∼2×10^6^ – 2×10^7^ cell mL^-1^ for co-cultivation with both bacterial strains) and cultured under photoautotrophic conditions. The bacterial cell density was determined by manual cell counting using hemocytometer or by determination of the colony-forming units (cfu) as a proxy for viable population density using the replica plating method (Jett et al., 1997).

The quantitative determination of the vitamin B_7_ (biotin) concentration in cell-free supernatants of the co-cultures and the axenic control was accomplished by using microbiological microtiter plate test (VitaFast, r-biopharm, Darmstadt, Germany) according to manufacturer instructions.

### Hydrocarbon extraction and analysis

Hydrocarbon extraction was performed according to the protocol published previously (Blifernez-Klassen et al., 2018). The dry extracted hydrocarbons were resuspended in 500 µL of *n*-hexane, containing an internal standard *n*-hexatriacontane (C_36_H_74_) and analysed *via* GC-MS and GC-FID as described before (Schwarzhans et al., 2014; Blifernez-Klassen et al., 2018).

### DNA sample collection and preparation

The samples from xenic *B. braunii* race A and B cultures (three biological replicates) were taken in the linear and stationary growth phases (after 9 and 24 days). Total community genomic DNA was extracted as previously described (Zhou et al., 1996). The quality of the DNA was assessed by gel electrophoresis and the quantity was estimated using the Quant-iT PicoGreen dsDNA Assay Kit (Invitrogen) and a Tecan Infinite 200 Microplate Reader (Tecan Deutschland GmbH).

### High-throughput 16S rDNA amplicon sequencing

To get insight into the community composition of *B. braunii* Race A and B, high-throughput 16S rDNA amplicon sequencing was performed as described recently by Maus et al (Maus et al., 2017). The primer pair Pro341F (5′-CCTACGGGGNBGCASCAG-3′) and Pro805R (5′-GACTACNVGGGTATCTAATCC-3′) was used to amplify the hypervariable regions V3 and V4 of diverse bacterial and archaeal 16S rRNAs (Takahashi et al., 2014). In addition, the primers also cover the 16S rDNA gene of algal chloroplasts and eukaryotic mitochondrial genomes. Furthermore, in a two PCR steps based approach, multiplex identifier (MID) tags and Illumina-specific sequencing adaptors were added to each amplicon. Only amplicons featuring a size of ∼460 bp were purified using AMPureXP® magnetic beads (Beckman Coulter GmbH, Brea, California, USA). Resulting amplicons were qualitatively and quantitatively analyzed using the Agilent 2100 Bioanalyzer system (Agilent Inc., Santa Clara, California, USA) and pooled in equimolar amounts for paired-end sequencing on the Illumina MiSeq system (Illumina, San Diego, California USA). This sequencing approach provided ∼150,000 reads per sample. An in-house pipeline as described previously (Wibberg et al., 2016) was used for adapter and primer trimming of all samples. For amplicon processing, a further in house amplicon processing pipeline including FLASH (Magoč and Salzberg, 2011), USEARCH (Edgar, 2010) (v8.1), UPARSE (Edgar, 2013) and the RDP (Wang et al., 2007) classifier (v2.9) was applied as described recently (Liebe et al., 2016; Klassen et al., 2017). In summary, unmerged sequences resulting by FLASH (Magoč and Salzberg, 2011) (default settings +-M 300) were directly filtered out. Furthermore, sequences with >1Ns (ambiguous bases) and expected errors >0.5 were also discarded. Resulting data was further processed and operational taxonomic units (OTUs) were clustered by applying USEARCH (Edgar, 2010) (v8.1). The resulting OTUs were taxonomically classified using the RDP (Wang et al., 2007) classifier (v2.9) in 16S modus (Threshold >0.8) and compared to the nt database by means of BLASTN (O’Leary et al., 2016). In the last step, raw sequence reads were mapped back onto the OTU sequences in order to get quantitative assignments.

### (Meta)Genome sequencing, assembly, binning and annotation

For each of the four samples (*B. braunii* race A and B from linear (T1) and stationary (T2) growth phase), the genomic DNA from three replicates was pooled. To obtain the metagenome sequence, four whole-genome-shotgun PCR-free sequencing libraries (Nextera DNA Sample Prep Kit; Illumina, Munich, Germany) were generated based on the manufacturer’s protocol representing different time points of the cultivation and different algae communities. The four libraries were sequenced in paired-end mode in a MiSeq run (2 x 300 bp) resulting in 11,963,558 and 11,567,552 reads for race A T1 and T2; 6,939,996 and 12,031,204 reads for race B T1 and T2, respectively (total 12.75 Gb). The *de novo* metagenome assembly was performed using the Ray Meta assembler (Boisvert et al., 2012) (v2.3.2) using a k-mer size of 41 and default settings. For contigs with length larger than 1 kb (n=127,914), the total assembly size was 338.5 Mb, the N50=2,884 and the largest contig 1.59 Mb. All processed raw sequence reads of the four samples were aligned to the assembled metagenome contigs by means of Bowtie 2 (Langmead and Salzberg, 2012) (v2.2.4). By using SAMtools (Li et al., 2009) (v1.6), the SAM files were converted to BAM files. These files were sorted and read mapping statistics were calculated. The following portions of reads could be assembled to the draft metagenomes: 70.1% and 77.5% for race A (T1 and T2); 51.0% and 62.9% for race B (T1 and T2). For binning of the metagenome assembled genomes (MAGs), the tool MetaBAT (v0.21.3) (Kang et al., 2015) was used with default settings. Completeness and contamination of the resulting MAGs were tested with BUSCO (Simão et al., 2015; Waterhouse et al., 2018) (v3.0.), using the Bacteria and Metazoa data set (Table S1).

For sequencing of the isolated species, 4 µg of purified chromosomal DNA was used to construct PCR-free sequencing library and sequenced applying the paired protocol on an Illumina MiSeq system, yielding ∼1.71 Mio Reads (517 Mb in 3 Scaffolds, 58 Contigs) for *Mycobacterium* sp. Bb-A and ∼1.70 Mio Reads (503 Mb in 6 Scaffolds, 65 Contigs) for *Pimelobacter* sp. Bb-B. Obtained sequences were *de novo* assembled using the GS *de novo* Assembler software (v2.8, Roche). An *in silico* gap closure approach was performed as described previously (Wibberg et al., 2011). MinION sequencing library with genomic DNA from *Brevundimonas* sp. Bb-A was prepared using the Nanopore Rapid DNA Sequencing kit (SQK-RAD04) according to the manufacturer’s instructions with the following changes: The entry DNA amount was increased to 800 ng and an AMPure XP bead cleanup was carried out after transposon fragmentation. Sequencing was performed on an Oxford Nanopore MinION Mk1b sequencer using a R9.5 flow cell, which was prepared according to the manufacturer’s instructions. MinKNOW (v1.13.1) was used to control the run using the 48h sequencing run protocol; base calling was performed offline using albacore (v2.3.1). The assembly was performed using canu v1.7 (Koren et al., 2017), resulting in a single, circular contig (170 Mb, 1kb to 17.5kb). This contig was then polished with Illumina short read data from the metagenome data set using pilon (Walker et al., 2014), run for sixteen iterative cycles. bwa-mem (Li and Wren, 2014) was used for read mapping in the first eight iterations and bowtie2 v2.3.2 (Langmead and Salzberg, 2012) in the second set of eight iterations. Annotation of the draft chromosomes was performed within the PROKKA (Seemann, 2014) (v1.11) pipeline and visualized using the GenDB 2.0 system (Meyer et al., 2003).

### Gene prediction, taxonomic assignment and functional characterization

Gene prediction on the assembled contigs of the MAGs was performed with Prodigal (Hyatt et al., 2010) (v2.6.1) using the metagenome mode (“-p meta”). All genes were annotated and analyzed with the in-house EMGB annotation system using the databases NCBI NR (O’Leary et al., 2016), Pfam (Finn et al., 2014) and KEGG (Kanehisa et al., 2017). Based on DIAMOND (Buchfink et al., 2015) (v0.8.36) hits against the NCBI-NR (O’Leary et al., 2016) database, all genes were subject to taxonomic assignment using MEGAN (Huson et al., 2016). Relationships to metabolic pathways were assigned based on DIAMOND (Buchfink et al., 2015) hits against KEGG (Kanehisa et al., 2017). For Pfam (Finn et al., 2014) annotations, pfamscan was used with default parameters. The genomes were manually inspected using an E-value cutoff of 1×10^−10^ for several specific sequences listed Additional file 2, Table S6. The profiling of the carbohydrate-active enzymes (CAZy) encoded by the *B. braunii* community members was accomplished using dbCAN2 (Zhang et al., 2018) metaserver (Table S7). The identification, annotation and analysis of the secondary metabolite biosynthesis gene clusters within the MAGs and complete genomes were accomplished *via* the antiSMASH (Blin et al., 2017) (v4.0) web server with default settings (Table S9).

### Phylogenetic analyses

To assign and phylogenetically classify the MAGs, a core genome tree was created by applying EDGAR (Blom et al., 2016) (v2.0). For calculation of a core-genome-based phylogenetic tree, the core genes of all MAGs and the three isolates *Mycobacterium* sp. Bb-A, *Brevundimonas* sp. Bb-A and *Pimelobacter* sp. Bb-B, corresponding to selected reference sequences were considered. The implemented version of FASTtree (Price et al., 2010) (using default settings) in EDGAR (Blom et al., 2016) (v2.0) was used to create a phylogenetic Maximum-Likelihood tree based on 53 core genes (see Table S4) of all genomes. Additionally, the MAGs were taxonomically classified on species level by calculation of average nucleic and amino acid identities (ANI, AAI) by applying EDGAR (Blom et al., 2016) (v2.0). For phylogenetic analysis of the 16S rDNA amplicon sequences, MEGA (Kumar et al., 2016) (v7.0.26) was used. Phylogenetic tree was constructed by means of Maximum-Likelihood Tree algorithm using Tamura 3-parameter model. The robustness of the inferred trees was evaluated by bootstrap (1000 replications).

## Availability of data and materials

The high-throughput 16S rDNA amplicon sequencing datasets obtained during the present work are available at the EMBL-EBI database under BioProjectID PRJEB21978. The metagenome datasets of *Botryococcus braunii* consortia as well as the resulting MAGs have been deposited under the BioProjectIDs PRJEB26344 and PRJEB26345, respectively. The genomes of the isolates *Brevundimonas* sp. Bb-A, *Mycobacterium* sp. BbA or *Pimelobacter* sp. Bb-B are accessible under BioProjectIDs PRJNA528993, PRJEB28031 und PRJEB28032, respectively. The communities most abundant ten high-quality draft as well as isolate genomes are also accessible via EMGB platform (https://emgb.cebitec.uni-bielefeld.de/Bbraunii-bacterial-consortium/).

## Authors’ contributions

O.B.K., V.K, and O.K. conceived this study. O.B.K., A.S., A.B., J.K. and O.K. supervised experiments and analyses. O.B.K., V.K., E.C. and S.C. accomplished all physiological studies. O.B.K., D.W., A.S., C.H., C.R., A.B., V.K., A.A.D., A.W., J.K., O.R., J.B. and A.G. performed the sequencing and the bioinformatic analyses. O.B.K., V.K., A.B. and O.K. wrote the original draft manuscript with contributions from all authors. O.B.K., V.K., A.B and O.K. reviewed and edited the manuscript. All authors read and approved the final manuscript.

## Acknowledgements

We are thankful to Joao Gouveia and Douwe van der Veen (Wageningen University, The Netherlands) for providing *Botryococcus braunii* race A and B (CCALA778 and AC761) stock cultures. Furthermore, we are grateful to the Center for Biotechnology (CeBiTec) at Bielefeld University for access to the Technology Platforms.

## Funding

This work was supported by the European Union Seventh Framework Programme (FP7/2007-2013) under grant agreement 311956 (relating to project “SPLASH – Sustainable PoLymers from Algae Sugars and Hydrocarbons”). The bioinformatics support by the BMBF-funded project “Bielefeld-Gießen Center for Microbial Bioinformatics” – BiGi (grant 031A533) within the German Network for Bioinformatics Infrastructure (de. NBI) is gratefully acknowledged.

## Ethics approval and consent to participate

Not applicable.

## Competing interests

The authors declare that they have no competing interests

## Additional files

### Supplementary Materials

Figure S1: Characteristics and taxonomic profiling of xenic *Botryococcus braunii* race A and B cultures. Illustrated are the typical colony-forming cells of *B. braunii* races A and B with a complex structure size of approximately >100µm in diameter. The table shows the mean values (three biological and technical replicates, SE; n=9) of algal content of biomass, hydrocarbons and carbohydrates at the sampling time points (linear and stationary growth phases). DW, culture dry weight.

Figure S2: Taxonomic profiling of *Botryococcus braunii* associating community. Relative abundance of detected taxa (phylum level) based on (**a**) high-throughput 16S rRNA gene amplicons and (**b**) metagenomic reads. The evaluation of 16S rDNA amplicons are detailed in Table S3. Metagenome datasets were taxonomically assigned via MEGAN (200,000 randomly subsampled reads, minimum 50 reads per taxonomic group). The relatively high proportion of unassigned reads likely results from the fact that the algal and some bacterial genomes are not disposed in the used NCBI-NR database. The analysis revealed the occurrence of the phyla Euryarchaeota and Viruses belonging to the family of *Microviridae* (bacteriophages with a single-stranded DNA genome).

Figure S3: Microbiome accompanying the *Botryococcus braunii* consortia. Illustration of the relative abundance and phylogenetic distribution of bacterial genera detected via high-throughput 16S rDNA amplicon sequencing approach (for details see Table S3).

Figure S4: Phylogenetic classification of selected *Botryococcus braunii* consortia members. Illustrated is the phylogenetic characterization of the high-quality draft and complete genomes of *B. braunii* accompanying bacterial community. Maximum-likelihood-based phylogenetic tree built out of a core of 53 conserved marker genes (Table S4) per analyzed genome (bold). Black, grey and white circles represent the nodes with calculated bootstrap values of 100, 99 and <75%, respectively (1,000 replicates).

Figure S5: Comparison and taxonomic assignment of different *Botryococcus braunii* metagenomes. Datasets obtained during the present study (races A and B, T1 and T2 (linear and stationary growth phases, respectively) were compared to the datasets of Guadeloupe race B strain (Condition A-C supplemented with citric acid as organic carbon source and vitamins (B_1_, B_7_ and B_12_), with A: initial consortium, B: washed culture, C: antibiotics-treated (ciprofloxacin)) (Sambles et al. 2017). For each sample, 200,000 metagenomic reads were randomly subsampled and subjected to the taxonomic assignment using DIAMOND and MEGAN (minimum 50 reads per taxonomic group). The circle size represents the amount of classified reads for the respective taxonomic group.

Figure S6: Hydrocarbon accumulation profiles of axenic and xenic *B. braunii* cultures. Shown is the hydrocarbon content observed in the samples of (**a**) axenic *B. braunii* (green) as well as in presence of (**b**) *Mycobacterium* sp. Bb-A (orange) and (**c**) *Brevundimonas* sp. Bb-A (blue) during the course of cultivation. The data serves as additional information for Figure 2.

Table S1: Features of the metagenome assembled genomes (MAGs). Assembly and annotation characteristics (*e.g.* Total size, N50, etc.) are listed according to the respective MAG. The completeness as well as the contamination degree of the MAGs was assessed via BUSCO (v3.0.). P- and E-Modi refer to prokaryotic MAGs and eukaryotic genome fragments, respectively. The high-quality MAGs were classified via single-copy marker genes, *e.g. rpoB, rpoD* etc., detected by BUSCO. The metagenomic datasets of *B. braunii* race A and B were mapped against the MAGs, and represented as percentage of mapped reads.

Table S2: Characteristics of high-quality draft and complete genomes. Summary of the taxonomy distribution and annotation features of ten high-quality MAGs and two complete genomes (completeness >80%, contamination <10%, Table S2) obtained from the *B. braunii* race A and B consortia. Shown are similarity mean values based on AAI (genome to genome comparison). Genomes with similarity values >95% are regarded as the same species as the reference. MAG, metagenome assembled genome; AAI, average amino acid identities; CDS, Coding DNA sequences.

#### Supplementary Discussion

Chapter 1 | Metagenomic survey of the *Botryococcus braunii* consortia

Chapter 2 | Elucidation of the genetic portfolio of the bacterial community

### Supplementary Tables. (XLSX, 330 KB)

Table S3: Statistics and taxonomic classification of 16S rRNA gene amplicon sequencing of the *B. braunii* consortia. Datasets include the analysis of the high-throughput 16S rDNA amplicon sequencing results and are supplemental information for Figures S2a and S3. Abbreviations: OTU, operational taxonomic unit; T, time point.

Table S4: List of conserved core genes. Dataset serves as supplemental information for Figure S4 and includes conserved marker (core) genes shared among the 31 genomes, used for the construction of the Maximum-likelihood-based (ML) phylogenetic tree.

Table S5: List of all detected functional pathways encoded in the high-quality draft and complete bacterial genomes. Datasets include KEGG-based functionally assigned pathways of the annotated high-quality MAGs and complete genomes imported to the EMGB platform. (elastic metagenome browser, https://emgb.cebitec.uni-bielefeld.de/Bbraunii-bacterial-consortium/). Displayed are the total number of the enzymes and the corresponding EC numbers encoded within the particular genome for the respective pathway. Bold marked pathway maps have been selected for Figure 1b.

Table S6: List of selected manually inspected genes within the high-quality draft and complete bacterial genomes. Datasets include manual BLASTp screening results (E-value threshold 1×10^−10^) of selected genes.

Table S7: List of Carbohydrate-active enzymes (CAZy) encoded by the *B. braunii* bacterial community members. Datasets include Carbohydrate-active enzymes (CAZy) encoded within the high-quality draft and complete bacterial genomes and serve as additional information for Figure 1b. The gene search within the high-quality draft and complete genomes was accomplished via dbCAN2 metaserver (Zhang et al. 2018).

Table S8: List of all detected B-vitamin biosynthesis genes encoded by the *B. braunii* bacterial community members. Datasets include the function description (EC number) as well as the corresponding genes of the key enzymes involved in the de novo biosynthesis of B-vitamins and serve as supplemental information for Figure 3a. The genes involved in the particular B-Vitamin pathway were searched (based on EMGB annotation system and manual BLASTp search) within the high-quality draft and complete bacterial genomes. √*, alternative gene identified via manual BLASTp search. Abbreviations: *M*. Bb-A, *Mycobacterium* sp. Bb-A; *B*. Bb-A, *Brevundimonas* sp. Bb-A; *P*.Bb-B, *Pimelobacter* sp. Bb-B.

Table S9: List of all detected secondary metabolite biosynthetic gene clusters. The datasets include the identified secondary metabolite biosynthetic gene clusters encoded by the high-quality draft and complete genomes of the *B. braunii* associates and serve as additional information for Figure 1b. The analysis was accomplished using the antibiotics & Secondary Metabolite Analysis Shell (antiSMASH, Version 4.1.0, Blin et al. 2017).

## Notes

#### Summary of Updates

The manuscript has been completely revised and contains new findings that support the hypotheses of the first version.

