## Supplementary Information for "Phytoplankton consortia as a blueprint for mutually beneficial eukaryote-bacteria ecosystems: Biocoenosis of *Botryococcus* consortia"

### Figure S1 |

#### Characteristics and taxonomic profiling of xenic *Botryococcus braunii* race A and B cultures

Illustrated are the typical colony-forming cells of *B. braunii* races A and B with a complex structure size of approximately >100µm in diameter. The table shows the mean values (three biological and technical replicates, SE; n=9) of algal content of biomass, hydrocarbons and carbohydrates at the sampling time points (linear and stationary growth phases). DW, culture dry weight.

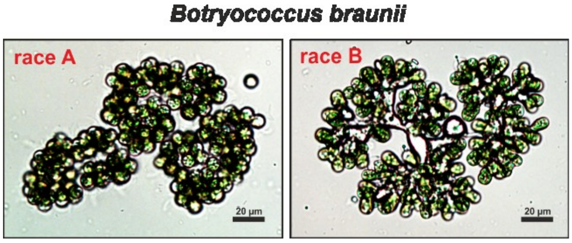


| | *B. braunii* race A | | *B. braunii* race B | |
|---|---|---|---|---|
| growth phase | linear T1 | stationary T2 | linear T1 | stationary T2 |
| biomass DW (g L$^{-1}$) | 1.1±0.1 | 3.6±0.4 | 1.1±0.1 | 2.7±0.2 |
| hydrocarbons (% of DW) | < 1% | < 1% | 59.3±3.6 | 57.4±8.3 |
| carbohydrates (% of DW) | 61.4±8.3 | 85.9±5.0 | 23.4±1.4 | 28.8±8.3 |

**Figure S2 |**

**Taxonomic profiling of *Botryococcus braunii* associating community**

Relative abundance of detected taxa (phylum level) based on (**a**) high-throughput 16S rRNA gene amplicons and (**b**) metagenomic reads. The evaluation of 16S rDNA amplicons are detailed in Table S3. Metagenome datasets were taxonomically assigned via MEGAN (Huson et al., 2016) (200,000 randomly subsampled reads, minimum 50 reads per taxonomic group). The relatively high proportion of unassigned metagenomic reads likely results from the fact that the algal and some bacterial genomes are not sequenced and disposed in the used NCBI-NR (O'Leary et al., 2016) database. The analysis revealed the occurrence of the phyla Euryarchaeota and Viruses belonging to the family of *Microviridae* (bacteriophages with a single-stranded DNA genome).

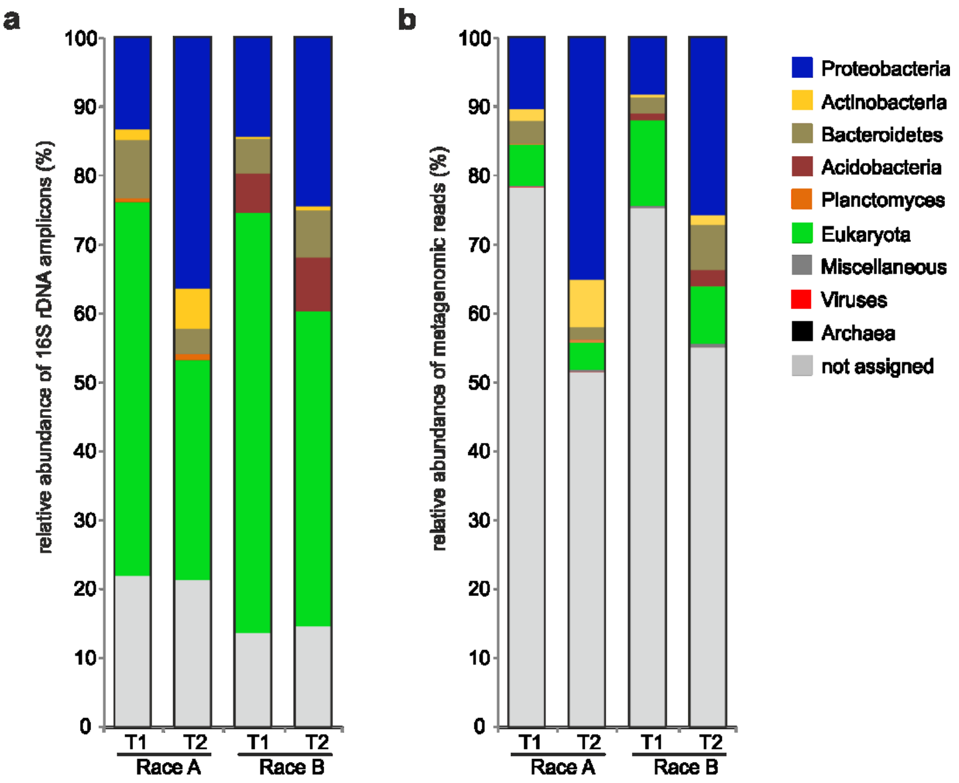

**Figure S3 |**

**Microbiome accompanying the *Botryococcus braunii* consortia**

Illustration of the relative abundance and phylogenetic distribution of bacterial genera detected via high-throughput 16S rDNA amplicon sequencing approach (for details see Table S3).

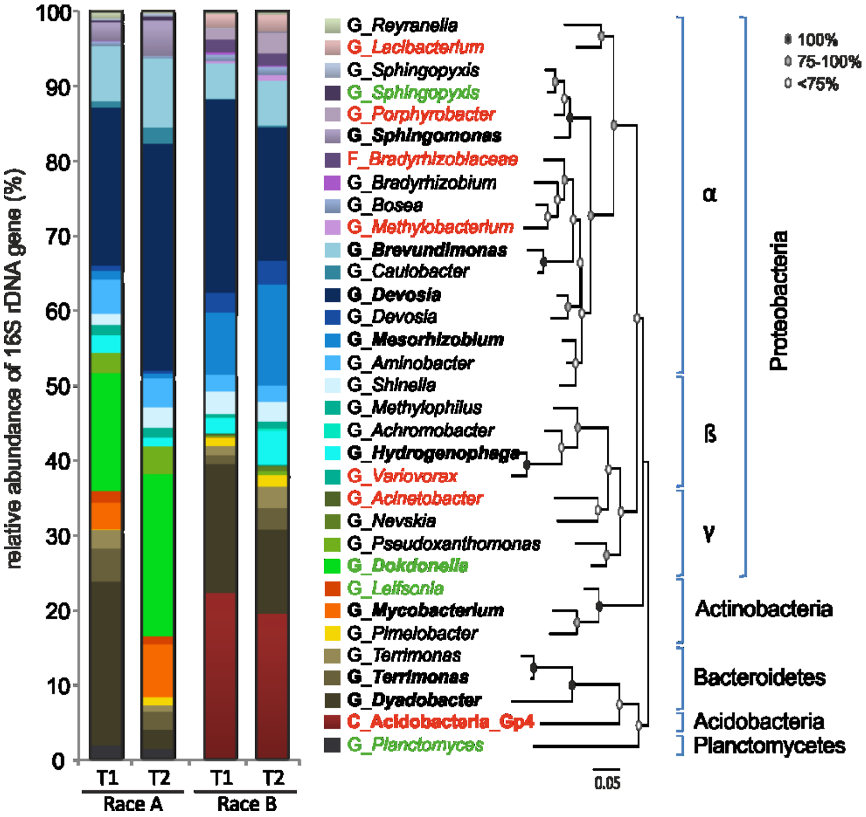

**Figure S4 |**

**Phylogenetic classification of selected *Botryococcus braunii* consortia members**

Illustrated is the phylogenetic characterization of the high-quality draft and complete genomes of *B. braunii* accompanying bacterial community. Maximum-likelihood-based phylogenetic tree built out of a core of 53 conserved marker genes (Table S4) per analyzed genome (bold). Black, grey and white circles represent the nodes with calculated bootstrap values of 100, 99 and <75%, respectively (1,000 replicates).

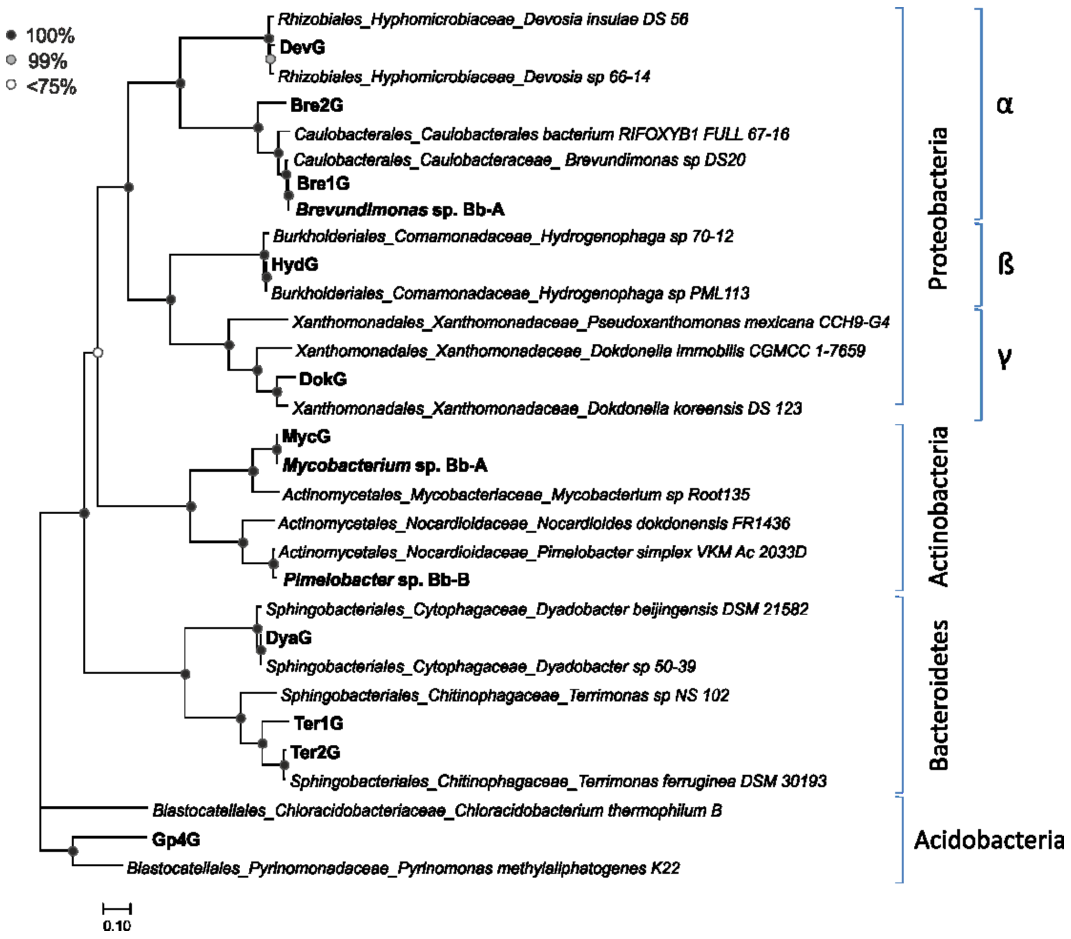

**Figure S5 |**

**Comparison and taxonomic assignment of different *Botryococcus braunii* metagenomes**

Datasets obtained during the present study (races A and B, T1 and T2 (linear and stationary growth phases, respectively)) were compared to the datasets of Guadeloupe race B strain (Condition A-C, supplemented with citric acid as organic carbon source and vitamins ($B_1$, $B_7$ and $B_{12}$); with A: initial consortium, B: washed culture, C: antibiotics-treated (ciprofloxacin)) (Sambles et al., 2017). For each sample, 200,000 metagenomic reads were randomly subsampled and subjected to the taxonomic assignment using DIAMOND (Buchfink et al., 2015) and MEGAN (Huson et al., 2016) (minimum 50 reads per taxonomic group). The circle size represents the amount of classified reads for the respective taxonomic group.

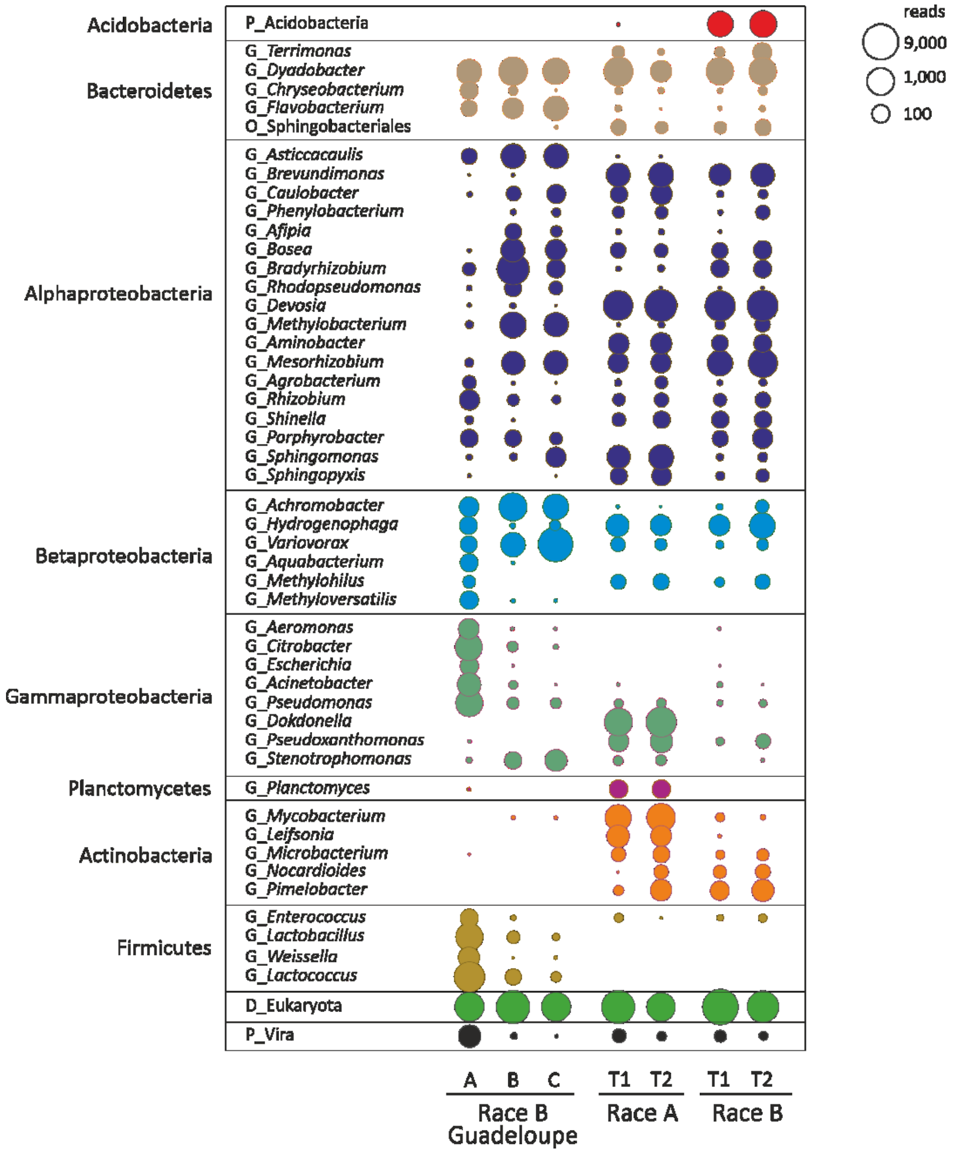


The re-processing of the *B. braunii* Guadeloupe strain metagenome data (Sambles et al., 2017) also shows data qualitatively but not quantitatively similar to the *Botryococcus* consortia analyzed in this study. The direct comparison of the different metagenome datasets revealed the coinciding presence of the individual genera (especially members of *Bacteroidetes*, *Alpha*- and *Betaproteobacteria*), although massively varying in the abundance pattern. The communities differ regarding the presence of the Gram-positive bacterial representatives: while the consortia obtained from our studies harbored the members of *Actinobacteria*, the members of the phyla *Firmicutes* were strongly represented in the Guadeloupe strain(Sambles et al., 2017). This evaluation of the *B. braunii* consortia under changing cultivation conditions has emphasized that depending on the physiological condition (*e.g.* supplementation of vitamins or antibiotics (Sambles et al., 2017)), the relative abundance of certain species may widely vary, suggesting the existence of many bacterial taxa with functional redundancy (Allison and Martiny, 2008).

**Figure S6 |**

**Hydrocarbon accumulation profiles of axenic and xenic *B. braunii* cultures**

Shown is the hydrocarbon content observed in the samples of (**a**) axenic *B. braunii* as well as in presence of (**b**) *Mycobacterium* sp. Bb-A (orange) and (**c**) *Brevundimonas* sp. Bb-A (blue) during the course of cultivation. The data serves as additional information for Figure 2.

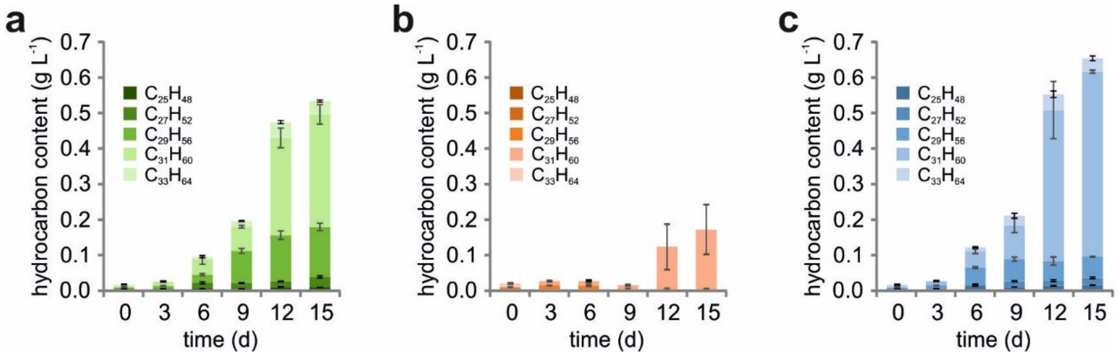

### Table S1 |

#### Features of the metagenome assembled genomes (MAGs).

Assembly and annotation characteristics (*e.g.* Total size, N50, etc.) are listed according to the respective MAG. The completeness as well as the contamination degree of the MAGs was assessed via BUSCO (Simão et al., 2015; Waterhouse et al., 2018) (v3.0.). P- and E-Modi refer to prokaryotic MAGs and eukaryotic genome fragments, respectively. The high-quality MAGs were classified via single-copy marker genes, *e.g. rpoB*, *rpoD* etc., detected by BUSCO (Simão et al., 2015; Waterhouse et al., 2018). The metagenomic datasets of *B. braunii* race A and B were mapped against the MAGs, and represented as percentage of mapped reads.

| | | MAG characteristics | | | | | BUSCO analysis | | | Mapping to the metagenomic reads | | | |
|---|---|---|---|---|---|---|---|---|---|---|---|---|---|
| | | | | | | | | | | *B. braunii* race A growth phase | | *B. braunii* race B growth phase | |
| MAG | Designation | Total size | N50 | Contig number | Largest Contig | Classification (Order) | Completeness | Duplicates | Modus | linear | stationary | linear | stationary |
| 1 | Dev-G | 5,369,495 | 947,618 | 20 | 1,587,146 | Rhizobiales | 94.60% | 0.70% | P | 2.73% | 13.17% | 3.17% | 5.14% |
| 2 | Dok-G | 4,393,516 | 994,039 | 7 | 1,352,042 | Xanthomonadales | 89.20% | 0.00% | P | 3.05% | 11.33% | 0.00% | 0.01% |
| 3 | Gp4-G | 4,376,278 | 1,138,557 | 5 | 1,579,402 | uc_Acidobacteria_Gp4 | 94.60% | 0.00% | P | 0.00% | 0.00% | 2.70% | 6.08% |
| 4 | Dya-G | 7,938,172 | 689,491 | 37 | 1,055,532 | Cytophagales | 91.90% | 4.10% | P | 2.28% | 0.74% | 1.86% | 3.42% |
| 5 | | 249,714 | 9,363 | 31 | 21,386 | uc_Trebouxiophyceae | 0.00% | 0.00% | E | 0.26% | 0.16% | 0.00% | 0.00% |
| 6 | | 331,094 | 7,900 | 47 | 19,793 | uc_Trebouxiophyceae | 0.00% | 0.00% | E | 0.40% | 0.24% | 0.00% | 0.00% |
| 7 | | 349,076 | 5,684 | 65 | 16,507 | uc_Trebouxiophyceae | 0.00% | 0.00% | E | 0.31% | 0.19% | 0.00% | 0.00% |
| 8 | | 457,400 | 7,931 | 66 | 17,190 | uc_Trebouxiophyceae | 0.70% | 0.00% | E | 0.35% | 0.22% | 0.00% | 0.00% |
| 9 | Ter1-G | 5,494,110 | 209,129 | 38 | 497,662 | Sphingobacteriales | 90.60% | 1.40% | P | 0.82% | 1.13% | 0.26% | 1.65% |
| 10 | Bre1-G | 3,248,831 | 226,467 | 25 | 389,267 | uc_Alphaproteobacteria | 96.60% | 0.00% | P | 0.94% | 2.72% | 0.03% | 0.10% |
| 11 | | 1,478,591 | 181,685 | 13 | 414,091 | Rhizobiales | 23.60% | 0.00% | P | 0.14% | 0.18% | 0.17% | 0.96% |
| 12 | | 240,645 | 68,507 | 5 | 89,489 | Burkholderiales | 10.80% | 0.00% | P | 0.02% | 0.04% | 0.02% | 0.13% |
| 13 | Myc-G | 6,010,666 | 365,564 | 25 | 949,347 | Corynebacteriales | 97.30% | 0.70% | P | 0.95% | 4.54% | 0.01% | 0.01% |
| 14 | Ter2-G | 4,576,776 | 18,593 | 328 | 75,155 | Sphingobacteriales | 81.80% | 0.70% | P | 0.34% | 0.16% | 0.27% | 1.63% |
| 15 | | 14,420,623 | 101,312 | 286 | 482,525 | Rhizobiales | 97.30% | 81.80% | P | 0.65% | 1.20% | 2.39% | 10.03% |
| 16 | Bre2-G | 3,095,339 | 105,164 | 52 | 214,209 | uc_Alphaproteobacteria | 92.60% | 8.10% | P | 0.04% | 0.11% | 0.68% | 1.99% |
| 17 | | 81,070,464 | 4,359 | 18,713 | 48,784 | uc_Trebouxiophyceae | 15.50% | 3.30% | E | 27.06% | 16.62% | 0.00% | 0.00% |
| 18 | | 12,107,853 | 3,243 | 3,647 | 10,635 | uc_Trebouxiophyceae | 1.30% | 0.00% | E | 0.00% | 0.00% | 5.04% | 3.29% |
| 19 | Hyd-G | 4,888,200 | 39,885 | 194 | 141,501 | Burkholderiales | 85.50% | 4.10% | P | 0.40% | 0.76% | 0.43% | 2.33% |
| 20 | | 1,393,646 | 3,624 | 389 | 8,353 | uc_Trebouxiophyceae | 0.30% | 0.00% | E | 0.00% | 0.00% | 0.44% | 0.25% |
| 21 | | 8,830,629 | 7,594 | 1,353 | 44,799 | Rhizobiales | 82.40% | 23.60% | P | 0.91% | 2.77% | 0.31% | 0.83% |
| 22 | | 2,026,801 | 3,533 | 549 | 17,353 | Propionibacteriales | 24.30% | 0.00% | P | 0.06% | 0.45% | 0.11% | 0.39% |
| 23 | | 774,780 | 3,607 | 209 | 18,620 | Micrococcales | 26.40% | 0.00% | P | 0.13% | 0.22% | 0.00% | 0.00% |
| 24 | | 1,040,113 | 4,083 | 259 | 14,226 | Sphingomonadales | 9.50% | 0.00% | P | 0.11% | 0.41% | 0.00% | 0.01% |
| 25 | | 1,788,866 | 4,030 | 440 | 13,282 | Xanthomonadales | 35.80% | 0.00% | P | 0.16% | 0.60% | 0.01% | 0.07% |
| 26 | | 553,426 | 3,237 | 152 | 28,072 | Sphingomonadales | 13.50% | 0.00% | P | 0.00% | 0.00% | 0.04% | 0.21% |
| 27 | | 251,679 | 3,077 | 79 | 6,433 | uc_Trebouxiophyceae | 1.30% | 0.00% | E | 0.00% | 0.00% | 0.02% | 0.07% |

### Table S2 |

#### Characteristics of high-quality draft and complete genomes

Summary of the taxonomy distribution and annotation features of ten high-quality MAGs and two complete genomes (completeness >80%, contamination <10%, Table S1) obtained from the *B. braunii* race A and B consortia. Shown are similarity mean values based on AAI (genome to genome comparison). Genomes with similarity values >95% are regarded as the same species as the reference. MAG, metagenome assembled genome; AAI, average amino acid identities; CDS, Coding DNA sequences.

| | Total lengh (Mb) | CDS number | GC content (%) | Class | Order | Close relative | | |
|---|---|---|---|---|---|---|---|---|
| | | | | | | % Similarity | Species | Accession |
| **DevG** | 5.369 | 5226 | 66.01 | Alpha-proteobacteria | Rhizobiales | 93.55 | *Devosia* sp. 66-14 | MKVB00000000 |
| **DokG** | 4.394 | 3582 | 70.14 | Gamma-proteobacteria | Xanthomonadales | 85.56 | *Dokdonella koreensis* DS-123 | CP015249 |
| **Gp4G** | 4.376 | 4155 | 52.5 | Blastocatellia | Blastocatellales | 62.12 | *Pyrinomonas methylaliphatogenes* K22 | CBXV00000000 |
| **DyaG** | 7.939 | 6743 | 50.68 | Cytophagia | Cytophagales | 99.47 | *Dyadobacter* sp. 50-39 | MKSR00000000 |
| **Ter1G** | 5.495 | 4633 | 42.1 | Chitinophagia | Chitinophagales | 70.94 | *Terrimonas ferruginea* DSM 30193 | AUDS01000038 |
| **Ter2G** | 4.581 | 4155 | 47.88 | Chitinophagia | Chitinophagales | 97.7 | | |
| **Bre1G** | 3.249 | 3165 | 67.46 | Alpha-proteobacteria | Caulobacterales | 99.25 | *Brevundimonas* sp. DS20 | CP012897 |
| ***Brevundimonas* sp. Bb-A** | 3.282 | 3171 | 67.4 | Alpha-proteobacteria | Caulobacterales | 99.25 | | |
| **Bre2G** | 3.096 | 3130 | 67.45 | Alpha-proteobacteria | Caulobacterales | 73.75 | | |
| **HydG** | 4.891 | 4720 | 70.02 | Beta-proteobacteria | Burkholderiales | 99.54 | *Hydrogenophaga* sp. PML113 | MIYM01000009 |
| **MycG** | 6.011 | 5762 | 67.37 | Actinobacteria | Corynebacteriales | 73.85 | *Mycobacterium* sp. Root135 | LMEZ00000000 |
| ***Mycobacterium sp. Bb-A*** | 6.015 | 5739 | 67.37 | Actinobacteria | Corynebacteriales | 73.85 | | |
| ***Pimelobacter sp. Bb-B*** | 5.86 | 5713 | 72.51 | Actinobacteria | Corynebacteriales | 98.5 | *Pimelobacter simplex* VKM Ac-2033D | CP009896 |

### Supplementary Discussion

#### Chapter 1 | Metagenomic survey of the *Botryococcus braunii* consortia

We profiled four different communities using high-throughput 16S rDNA gene amplicon data and added 12.75 Gb of paired-end metagenome data. The analysis of the 16S rDNA amplicons and a read-based metagenome approach revealed that each of the *B. braunii* consortia consists of a single algal and various bacterial species as well as traces of Archaea and viruses (Figure S2). Each community is comprised of four bacterial phyla, consisting of at least 25 species with the members of the phylum *Proteobacteria* being the most prominent in both *B. braunii* races. Of 33 different bacterial genera, four were specific to race A, seven were specific to race B and 22 were shared (Figure S3). The relative abundance of plastid 16S rDNA sequences accounted up to 49 and 59% of all detected amplicon reads in race A and B samples in the linear growth phase and declined to approximately 26% and 44% at stationary phase, respectively (Figure S2a). At the same time, the 16S rDNA contribution of the accompanying microbial community increased. The community members varied quantitatively based on both race and time of sampling, but not qualitatively.

Both *B. braunii* races were dominated by the genera *Devosia* and *Dyadobacter* (up to 26% and 22% of the bacterial community) in the logarithmic growth phase, followed by *Brevundimonas* (7.4%), *Dokdonella* in race A (16%) and *Acidobacteria* in race B (22%) (Figure S3). During the cultivation, the relative abundancy of the dominant species in the race A consortium such as *Devosia* and *Dokdonella* increased at the expense of *Dyadobacter* in comparison to the linear growth phase. In the race B consortium, less abundant taxa such as *Mesorhizobium, Brevundimonas* and *Hydrogenophaga* increased at the expense of *Dyadobacter* and *Devosia*. The detected relative levels of both actinobacterial isolates were comparatively low (Figures S2 and S3). *Mycobacterium* isolate, obtained from the *B. braunii* race A community, was almost exclusively propagated therein and increased during stationary growth to 7% of all detected bacterial amplicons (Figure S3). *Pimelobacter* was isolated from the race B consortium, and is present in both communities in equally (≤1%) low abundances which increased slightly during stationary phase.

To test the relationships between *B. braunii* and the associative consortium, we explored the genetic potential of the abundant representatives of algal-bacterial consortia. We combined metagenome *de novo* assembly and differential coverage and tetra-nucleotide signature based binning of the *B. braunii* race A and B samples. This approach resulted in the reconstruction of ten high-quality (completeness >80%, contamination <10%) and ten fragmentary metagenome assembled genomes (MAGs) as well as seven eukaryotic genome fragments (for details see

Methods; Table S1). To identify taxa representing the ten high-quality draft MAGs, we constructed a phylogenetic tree built out of a core of 53 highly conserved marker genes per genome (Figure S4). All abundant taxa shared by both races were recovered, including the most abundant alphaproteobacterial MAG of the order *Rhizobiales*, designated DevG (similar to *Devosia* sp. 66-14, Table S2 and Figure S4). Additionally, MAGs of *Dyadobacter* and *Hydrogenophaga* were constructed (DyaG and HydG, similar to *Dyadobacter* sp. 50-39 and *Hydrogenophaga* sp. PML113, respectively). Another two alphaproteobacterial MAGs Bre1G and Bre2G of the order *Caulobacteriales* (similar to *Brevundimonas* sp. DS20), clustered together within the 16S rDNA amplicon approach (Figure S3) but were separated through the binning procedure (Table S1). We reconstructed sphingobacterial MAGs of the order *Chitinophagales*, Ter1G and Ter2G similar to different extent to *Terrimonas ferruginea* DSM30193. Further two MAGs, basically occurring in race A strain, included a gammaproteobacterial bacterium similar to *Dokdonella koreensis* DS-123 (DokG) and a member of *Corynebacteriales* similar to *Mycobacterium* sp. Root135 (MycG). An exclusively in *B. braunii* race B occurring member of the subdivision IV of the phyla *Acidobacteria* (Gp4G) is similar to *Pyrinomonas methylaliphatogenes* K22.

We also sequenced the bacterial isolates, *Pimelobacter* sp. Bb-B, *Mycobacterium* sp. Bb-A as well as *Brevundimonas* sp. Bb-A, and used the latter two for the co-cultivation experiments (Figures 2 and 4). After assembly and *in silico* finishing, both genomes were completed and represented by a single chromosome, each. The *Mycobacterium* sp. Bb-A and *Brevundimonas* sp. Bb-A genome sequences were identical to the MycG and Bre1G, respectively, indicating that both represent the same species (Table S2), thus underlining the quality of our metagenome assembly and binning approach. The *Pimelobacter* sp. Bb-B genome sequence resembled the MAG-22, which is only to 24% complete (Table S1).

**Chapter 2 | Elucidation of the genetic portfolio of the bacterial community**

We performed a KEGG-based quantitative functional assignment of the annotated high-quality MAGs and sequenced genomes by applying the elastic metagenome browser (EMGB, https://emgb.cebitec.uni-bielefeld.de/Bbraunii-bacterial-consortium/). Six of twelve genomes carried genes for bacterial chemotaxis and five of twelve genes for flagella synthesis, hinting at active motility and ability to move towards substrates in the medium (Figure 1b). *Botryococcus braunii* synthesizes aliphatic long-chain hydrocarbons, which accumulate in large quantities in the extracellular matrix of this species (Banerjee et al., 2002). Seven of twelve genomes carried gene homologs of already characterized hydrocarbon hydroxylases (alkB, ladA, almA and/or cytochrome P450 (CYP), Table S6) for the initial oxyfunctionalization of long-chain hydrocarbons (Rojo, 2010). While the most abundant consortia members like DevG and DyaG exhibited a very low enzyme number, the more rare *Mycobacterium* and *Pimelobacter* genomes contained a large portfolio of hydrocarbon monooxygenase/hydroxylase genes, and were the only genomes to carry non-heme membrane-associated monooxygenase alkB genes (Table S6).

Mucilaginous (exo-)polysaccharides as part of the internal fibrillar layer of the *B. braunii* cell wall represent another prospective carbon source for the bacterial community, consisting mainly of galactose but also of fucose and rhamnose (Banerjee et al., 2002; Blifernez-Klassen et al., 2018). The profiling of the carbohydrate-active enzymes (CAZymes (Lombard et al., 2014)), encoded by the members of *B. braunii* communities using dbCAN2 (Zhang et al., 2018) metaserver, revealed the presence of several putative enzyme families involved in the complex carbohydrate metabolism (Table S7). Eight out of twelve genomes coded for several carbohydrases for the degradation of poly- and oligosaccharides, generally present in the cell walls of microalgae, and contained genes for chitin disintegration and assimilation (Additional file 2, Table S7). However, six species showed a clear prevalence in the number of enzymes involved in the biosynthesis of carbohydrates (glycosyltransferases) compared to degradation (glycoside hydrolases). Some species like the members of *Bacteroidetes* contained a large number of CAZyme-encoding genes in their genomes (up to 5.9% of total gene content) with a clear prevalence to hydrolytic activity of complex carbohydrates (Table S7). For instance, the DyaG genome contains more genes encoding for the glycoside hydrolase (GH) families involved in the degradation of complex polymers (cellulose, xylan, chitin), while others, such as the genome of DevG, harbor enzymes acting on oligosaccharides (Lombard et al., 2014).

Looking at the primary metabolism, all genomes encode genes for processes essential for aerobic respiration (Figure 1b; Table S5). Genes that enable pyruvate fermentation to lactate, acetate and ethanol are also present, showing a versatility of the community members to switch between aerobic and anaerobic metabolisms depending on the oxygen availability. From the perspective of a microalga, the associated bacteria can be providers of micro- and macronutrients but also competitors for limiting nutrients (Cole, 1982). Based on their observed generic potential, individual species appear to be involved in the acquisition and re-mineralization of macronutrients and in the storage of nitrogen and phosphorous (in form of cyanophycin and Poly-P, Table S6), which benefit the microalga. Eight of twelve genomes encode genes for the synthesis of cyanophycin granule polypeptides (CGPs) (Rehm, 2010), however only four contained genes for the degradation of this C- and N-storage compound (Table S6). Related to this, all genomes carry genes for the synthesis but not for the degradation of polyphosphate granules (Poly-P), a polymeric reserve with a significant role in the regulation of enzymatic activities, gene expression and stress adaptation processes (Kulakovskaya et al., 2012). Suggesting an adaptation to phosphorous and iron limitation, all genomes contained genes relevant for phosphate starvation (*pst* system) or siderophore biosynthesis (*fhu* system).
